# Inactive PARP1 causes embryonic lethality and genome instability in a dominant-negative manner

**DOI:** 10.1101/2023.05.23.542022

**Authors:** Zhengping Shao, Brian J. Lee, Hanwen Zhang, Xiaohui Lin, Chen Li, Wenxia Jiang, Napon Chirathivat, Steven Gershik, Michael M. Shen, Richard Baer, Shan Zha

**Affiliations:** Institute for Cancer Genetics, Department of Pathology and Cell Biology, College of Physicians and Surgeons, Columbia University, New York City, NY 10032; Departments of Medicine, Genetics and Development, Urology, and Systems Biology, Herbert Irving Comprehensive Cancer Center, Columbia University Medical Center, New York, NY 10032, USA; Division of Pediatric Oncology, Hematology and Stem Cell Transplantation, Department of Pediatrics, College of Physicians & Surgeons, Columbia University, New York City, NY 10032

**Keywords:** PARP1, E988A mutation, poly-ADP-ribosylation (PARylation)

## Abstract

PARP1 is recruited and activated by DNA strand breaks, catalyzing the generation of poly-ADP-ribose (PAR) chains from NAD+. PAR relaxes chromatin and recruits other DNA repair factors, including XRCC1 and DNA Ligase 3, to maintain genomic stability. Here we show that, in contrast to the normal development of Parp1-null mice, heterozygous expression of catalytically inactive Parp1 (E988A, *Parp1^+/A^*) acts in a dominant-negative manner to disrupt murine embryogenesis. As such, all the surviving F1 *Parp1^+/A^* mice are chimeras with mixed *Parp1^+/AN^* (neoR retention) cells that act similarly to *Parp1^+/-^*. Pure F2 *Parp1^+/A^* embryos were found at Mendelian ratios at the E3.5 blastocyst stage but died before E9.5. Compared to *Parp1^-/-^* cells, genotype and expression-validated pure *Parp1^+/A^* cells retain significant ADP-ribosylation and PARylation activities but accumulate markedly higher levels of sister chromatin exchange and mitotic bridges. Despite proficiency for homologous recombination and non-homologous end-joining measured by reporter assays and supported by normal lymphocyte and germ cell development, *Parp1^+/A^* cells are hypersensitive to base damages, radiation, and Topoisomerase I and II inhibition. The sensitivity of *Parp1^+/A^* cells to base damages and Topo inhibitors in particular exceed *Parp1^-/-^* controls. The findings show that the enzymatically inactive PARP1 protein has a dominant negative role and establishes a clear physiological difference between PARP1 inactivation vs. deletion. As a result, the enzymatically inactive PARP1 has a much more deteriorating impact on normal tissues than previously estimated, providing a mechanism for the on-target side effect of PARP inhibitors used for cancer therapy.

**Significance Statement:** PARP1 is the primary target of PARP enzymatic inhibitors. The use of PARP inhibitors for cancer therapy is based not only on the extreme sensitivity of BRCA1/2-deficient cancer cells to PARP1 inhibition but also on the nonessential role of PARP1 in normal tissues. Here we show that in contrast to the normal development of Parp1-null mice, the mouse model expressing the catalytically inactive Parp1 on only one allele (E988A, *Parp1^+/A^*) dies embryonically with high levels of genomic instability. The results reveal the severe dominant-negative impact of catalytically inactive PARP1, indicating the presence of enzymatically inactive PARP1 is much more damaging to normal tissues than previously anticipated. These findings provide a mechanism for clinical PARP inhibitors’ unexpected normal tissue toxicity.

## Introduction

PARP1 belongs to the family of poly-ADP ribose polymerases (PARPs) that share a conserved ADP-ribose (ADPr) transferase (ART) domain. PARP1 and the related PARP2 are recruited to DNA strand breaks, where they are activated to catalyze formation of poly-ADP-ribose (PAR). PAR promotes chromatin relaxation and recruits other repair proteins, including the XRCC1-Ligase3 complex that ligates single-strand DNA nicks, such as those generated during base excision repair (BER)(1–3). PARP1 is more abundant than PARP2, has a higher affinity for diverse DNA lesions, and accounts for >80% of DNA damage-induced PARP activity(4). Yet, *Parp1*-null mice are viable, fertile, and of standard size(5). Indeed, the use of PARP enzymatic inhibitors (PARPi) for cancer therapy is based not only on the extreme sensitivity of BRCA1-or BRCA2-deficient cancer cells to PARP1 loss, but also on the nonessential role of PARP1 in normal tissues(6, 7).

In addition to blocking its enzymatic activity, PARPi also induces a phenotype termed “PARP trapping”, characterized by the persistence of damage-induced PARP1/2 foci and prolonged retention of PARP1/2 on damaged chromatin (8–11). Loss of PARP1, but not PARP2, causes marked resistance to PARP inhibitors (10, 11), highlighting the unique importance of PARP1 inhibition in cancer therapy. Several mechanisms have been proposed to explain PARP1 trapping (12). *In vitro,* PARPi prevents PARP1 auto-PARylation, which correlates with the release of PARP1 from DNA ends (9). But this mechanism alone cannot explain why PARPi with similar IC50s for enzymatic inhibition have different abilities to “trap” PARP1. Structural analyses later showed that the non-hydrolysable NAD+ analog benzamide adenine dinucleotide (BAD) could reverse-allosterically enhance PARP1 affinity for DNA ends (13), providing an explanation for the differential trapping by inhibitors with distinct chemical structures. PARPi attenuates PAR, which recruits XRCC1-LIG3 and other repair complexes (11). Using quantitative live cell imaging, we recently showed that PARP1 molecules exchange rapidly at micro-irradiation sites in cells (11), even in the presence of clinically-effective PARPi, suggesting that the PARPi-induced persistence of PARP1 foci may reflect continual recruitment of different PARP1 molecules to the unrepaired DNA lesion due to delayed repair. In this regard, the release and exchange of PARP1 from model DNA ends *in vitro* can also occur independently of auto-PARylation and in the presence of clinically relevant PARPi(12, 14–16).

Although the therapeutic effects of PARP inhibition and trapping on tumor cells have been studied extensively, less is known about the impact of inactive PARP1 on normal cells, despite accumulating evidence of clinical PARPi toxicities in both patients and inhibitor-treated wild-type mouse models (17–20). To address this question and understand the physiological impact of inactive PARP1 protein, we introduced a catalytically inactive point mutation into the endogenous Parp1 locus. Here we show that, in contrast to the normal development of Parp1-null mice, heterozygous expression of catalytically inactive Parp1 (E988A, *Parp1^+/A^*) acts in a dominant-negative manner to disrupt embryogenesis and genome stability. *Parp1^+/A^* cells are characterized by the elevated formation of mitotic bridges and sister chromatid exchanges. The genome toxicity is due to the accumulation of inactive PARP1 protein at DNA damage sites, but not the loss of ADP-ribosylation and PARylation activity. Mechanistically, we show that inactive PARP1-E988A is recruited to DNA damage sites where it forms persistent foci. Especially, the inactive Parp1 blocks the resolution of DNA damage induced by alkylating agents and topoisomerase inhibitors beyond the loss of PARP1, but not clean DNA double-strand break repair via homologous recombination (HR) or non-homologous end-joining (NHEJ), measured by lymphocyte development, meiosis, and reporter assays. The results demonstrate a clear difference between PARP1 inhibition vs PARP1 deletion on normal tissues, providing the molecular mechanism for the initially unexpected and significant adverse effects of PARPi.

## Results

### The generation and characterization of catalytically inactive Parp1 murine model

To address the impact of inactive PARP1 on normal issue, we generated the *Parp1^AN^* allele by introducing the catalytically inactive E988A missense mutation(21) and an FRT-NeoR-FRT cassette into the endogenous Parp1 locus of mice (Fig. 1A and 1B). The E988A mutation abrogates the PARylation activity of PARP1 without affecting its DNA binding(21). Since the FRT-NeoR-FRT cassette blocks transcription (Fig. 1A), *Parp1^AN^* behaves like a null allele and, as anticipated, *Parp1^+/AN^* pups were born at the expected Mendelian ratio (Fig.1C). To remove the cassette and generate the desired *Parp1^A^* allele, *Parp1^+/AN^* mice were bred with *Rosa^FLIP/FLIP^* mice that constitutively express FLIPase (Jax Strain No. 003946 in 129Sv). Surprisingly, the F1 *Parp1^+/A^* pups were significantly under-represented among the progeny (1/6 of expected) (Fig.1D). They were also ∼25% smaller than their littermates (Fig.1E) and many (∼45.8%) had kinked tails (Fig. 1F), a phenotype previously seen in genomic instability mouse models (*e.g.*, *H2ax*^-/-^ or *Brca1* mutants), but not in *Parp1^-/-^* mice, suggesting that catalytically inactive Parp1 polypeptides can disrupt genome integrity in a dominant-negative manner.

**Figure 1.**
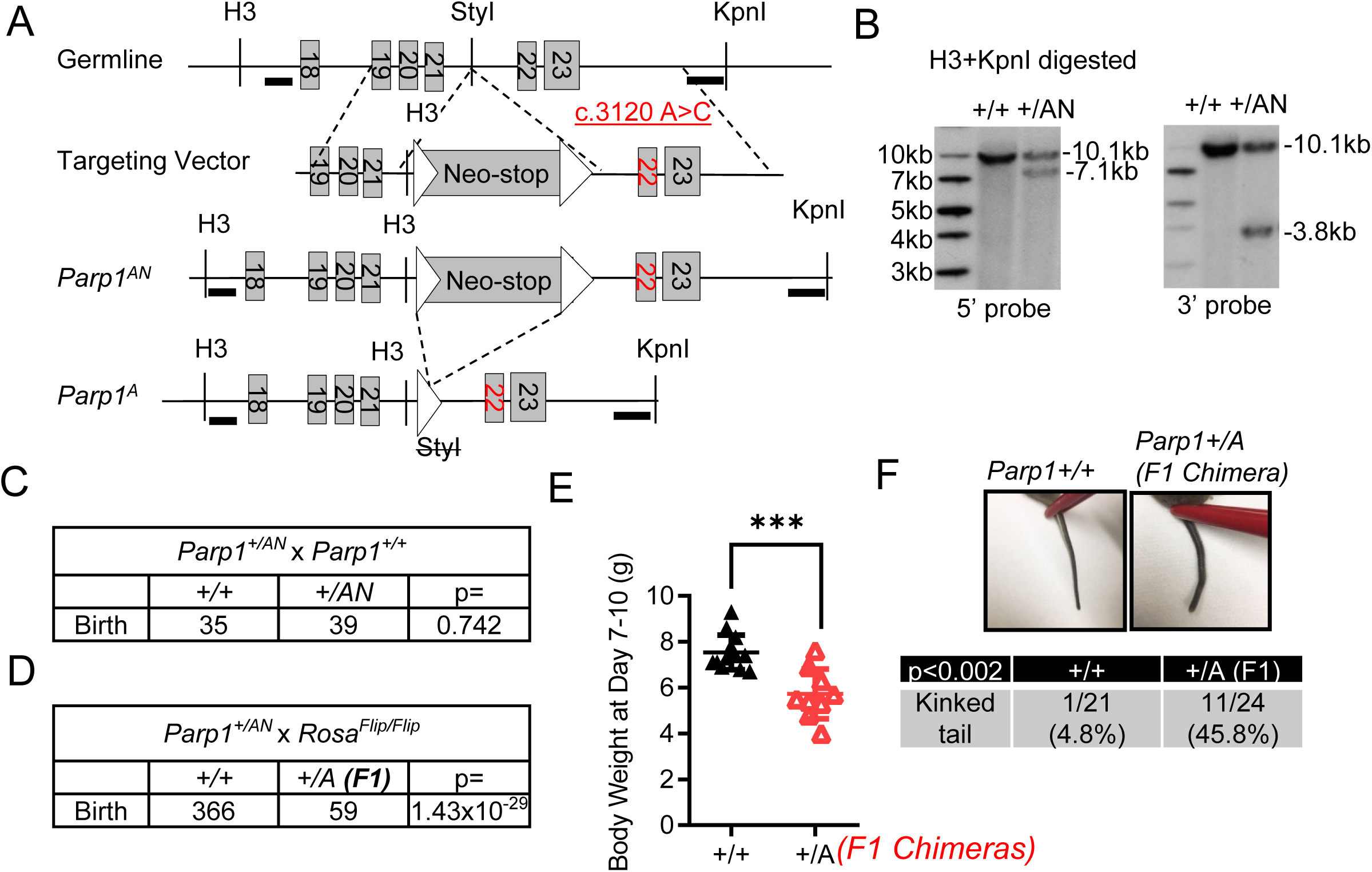
Expression of catalytically inactive Parp1 impairs murine development. (A) Mouse Parp1 E988 E->A targeting strategy. The schematic diagram represents the murine Parp1 germline structure (top row), targeting vector (second row), targeted allele (Parp1^AN^, third row), and the recombined E988A expression allele without neo (Parp1^A^, bottom row). The presence of the NeoR in the Parp1^AN^ allele interferes with the expression of Parp1, resulting in no or little Parp1 expression. Map not shown to scale. (B) Southern blot analysis of targeted *Parp1^+/AN^* ES cell DNA using 5’ and 3’ probes. Left: wild-type ES cells. Right: targeted ES cells. Top band: germline. Lower band: targeted. (C) The birth rate of F1 pups from *Parp1^+/AN^* x *Parp1^+/+^* crossing. P-values were calculated via the χ^2^ test. (D) The birth rate of F1 pups from *Parp1^+/AN^* x *Rosa^Flip/Flip^* crossing. P-values were calculated via the χ^2^ test. (E) Body weight of *Rosa^+/Flip^ Parp1^+/A^* and *Rosa^+/Flip^ Parp1^+/+^* pups. The student’s T-test calculated the p-value. ***: P<0.001. (F) Representative images and frequency of F1 *Parp1^+/A^* chimeras or *Parp1^+/+^* mice with kinked tails.

### F2 *Parp1^+/A^* mice die during early embryonic development

Curiously, all the F2 pups (n=52) of mating between *F1 Parp1^+/A^* mice and WT partners (both genders) were *Parp1^+/+^*, regardless of p53 status (Fig. 2A, p<3.9×10^-9^) and despite equal representation of *Parp1^E988A^* and *Parp1^+^* alleles in the sperm of F1 *Parp1^+/A^* mice (Fig. 2B). Timed mating showed that F2 *Parp1^+/A^* embryos were recovered with normal early-stage blastocyst morphology at E3.5 (Fig. 2C) but could not be recovered at E9.5, indicating F2 *Parp1^+/A^* animals likely suffered early embryonic lethality (Fig. 2A). Moreover, despite the known requirement for HR in germ cell development and meiosis, testis architecture (Fig. 2D), sperm counts (Fig. 2E), and sperm mobility were normal in young F1 *Parp1^+/A^* mice used for breeding (*SI Appendix*, Fig. S1A). The lack of F2 *Parp^+/A^* mice was perplexing given the viability of F1 *Parp1^+/A^* pups (Fig. 1D). We hypothesized that the surviving F1 *Parp1^+/A^* mice might be chimeras, which carry *Parp1^+/AN^* cells that had escaped FLIPase excision and whose presence, although low in frequency, support the embryogenesis of F1 *Parp1^+/A^* mice. To test this possibility, we re-genotyped F1 *Parp1^+/A^* mice by PCR using both the standard triplex primers (detecting AN, A, and + alleles) as well as primers that only detect the AN allele (*Fig. S1B*). All F1 *Parp1^+/A^* mouse tail DNAs tested AN-positive (allele frequency: 1-10%) in the AN-specific PCR assay (Fig. 2F-G and *Fig. S1C-D*). We note that the PCR product for the A allele shares 230bp sequence identical to that of the + allele, except the ∼80bp insertion corresponding to the FRT site and its surrounding sequence (*Fig. S1B*). As a result, when both A and + products are present (*e.g.,* in *Parp1^+/A^*), the two PCR products sometimes form a heteroduplex that migrates between the WT and A PCR product (Fig. 2F). Moreover, qPCR analyses of peripheral blood from F1 *Parp1^+/A^* mice at different ages showed a progressive increase of AN allele frequency from ∼1% at three weeks to ∼100% at 64 weeks (Fig. 2H). This is consistent with the low chimerism in the tail DNA collected at 7 days after birth (Fig. 2F-G and *Fig. S1C-D*). Thus, we concluded that pure heterozygosity abrogates the embryonic development of *Parp1^+/A^* mice.

**Figure 2.**
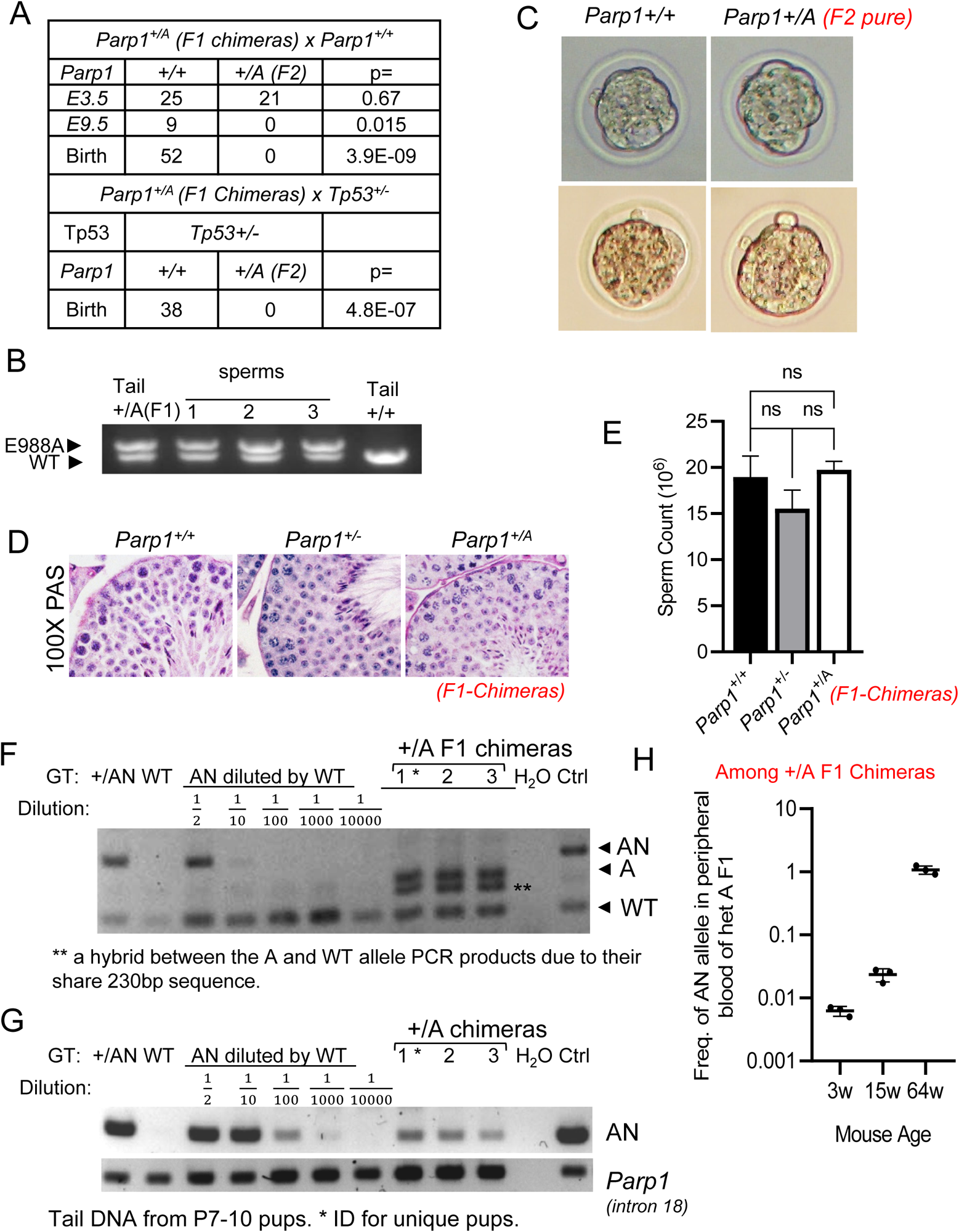
Heterozygous expression of inactive Parp1-E988A causes embryonic lethality in mice. (A) The frequency of F1 *Parp1^+/A^* pups from *Parp1^+/A^* x *Parp1^+/+^*crossing with or without *Tp53* deficiency at different developmental stages. P-values were calculated via the χ^2^ test. (B) PCR identification of *Parp1^A^* allele in F1 *Parp1^+/A^* mice sperm. (C) Morphology of F2 *Parp1^+/+^* and *Parp1^+/A^*early-stage blastocysts at E3.5. (D) Histology with H&E staining showing spermatogenesis in young (<8 weeks) male *Parp1^+/+^, Parp1^+/-^,* and *Parp1^+/A^*testes. (E) Epididymis mature sperm count of *Parp1^+/+^, Parp1^+/-^,* and *Parp1^+/A^* adult males. The P-value was calculated based on one-way ANOVA. ns: p>0.05. (F and G) Standard triplex tail PCR (F) and targeted AN-specific PCR (G) of F1 *Parp1^+/A^* mice and controls. (H) qPCR of *Parp1^AN^*in peripheral blood of F1 *Parp1^+/A^* mice of different ages.

### Normal lymphocyte development and spermatogenesis in viable young (<8 weeks) F1 *Parp1^+/A^* chimeric mice

PARP1 is activated by DNA breaks. To understand the mechanism of growth retardation, we then asked whether inactive Parp1 blocks DNA double-strand break (DSB) repair. Mammalian cells have two major DSB repair pathways –NHEJ or HR. Specifically, lymphocyte development requires the ordered assembly and subsequent modification of immunoglobulin (Ig) genes through two programmed DSB events - V(D)J recombination and Immunoglobulin Class Switch recombination (CSR) (22). Defects in the NHEJ pathway abrogate V(D)J recombination and significantly impair CSR (22), serving as a physiological readout of NHEJ. The *Parp1^+/AN^* chimerism in the F1 *Parp1^+/A^* mice is consistently <10% in peripheral blood by 15 weeks of age. So, we focused our analyses of the F1 Chimeric *Parp1^+/A^* on young mice only. Like in Parp1 null mice (5), B and T lymphocyte development measured by both relative frequency of immature and mature cells and developmental state-specific cellularity, were both normal in young F1 *Parp1^+/A^* mice (<8 weeks) (Fig. 3A-C). Furthermore, IgH CSR, a sensitivity measure for NHEJ (23), was also normal in the splenocytes purified from young F1 *Parp1^+/A^* mice (<8 weeks) (Fig. 3A, 3D). Since CSR efficiency can be influenced by cell proliferation, we further measured CSR by gating cells with the same cell division. No measurable defects in CSR were noted in *Parp1^+/A^* splenocytes even after being controlled for cell division (*Fig. S2A*). Given the normal spermatogenesis, a process that requires HR, in young *Parp1^+/A^* chimeras, these data together suggest that heterozygous expression of inactive PARP1 does not have a major impact on NHEJ or HR. However, due to the chimerism, although low, we would not be able to rule out minor impacts on HR and NHEJ in the *Parp1^+/A^* mice based on the apparent normal development in F1 chimeric *Parp1^+/A^* mice.

**Figure 3.**
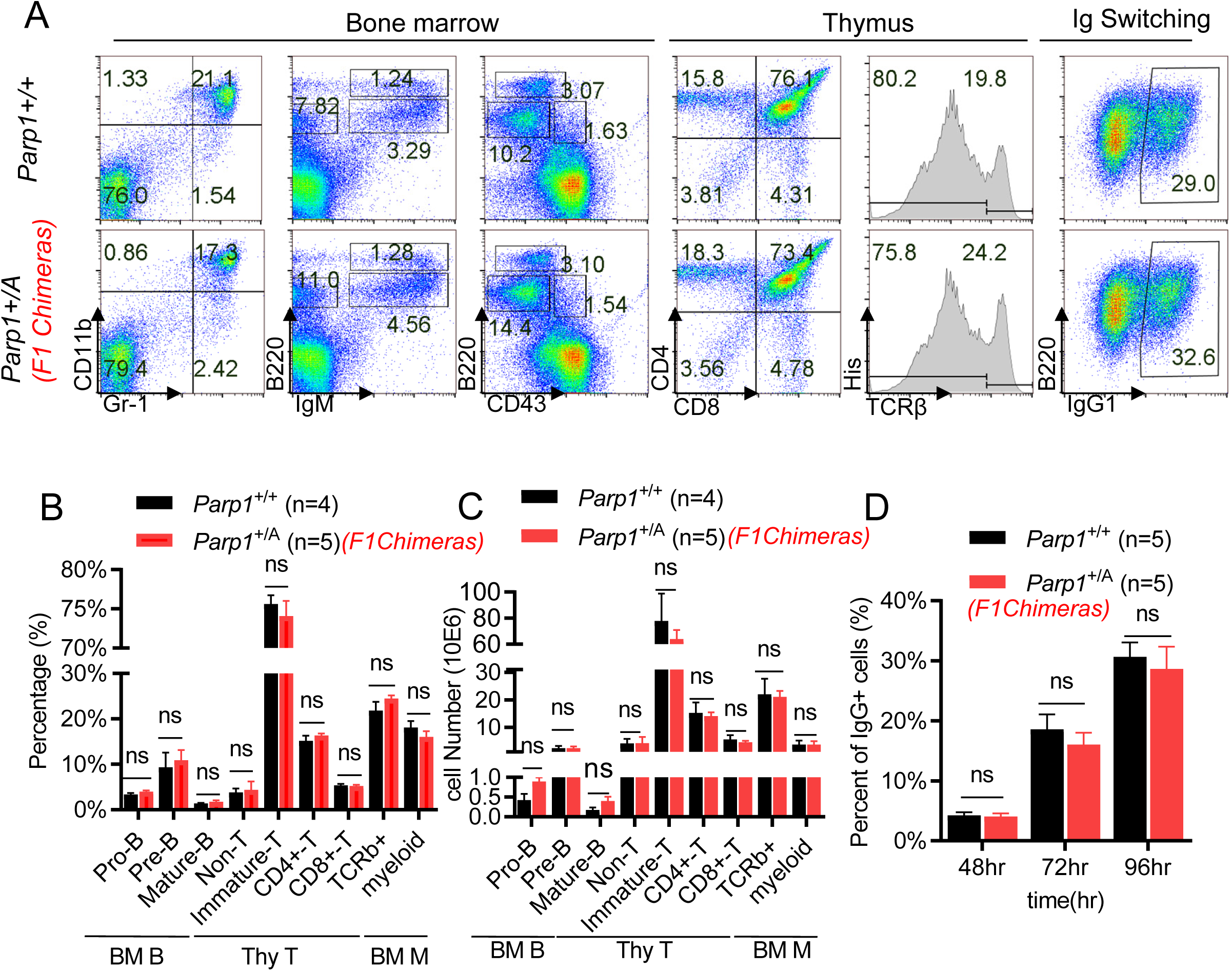
Young F1 chimeric *Parp1^+/A^* mice have normal lymphocyte development. (A-D) Representative flow cytometry analyses (A), relative percentage (B), and absolute cell count (C) of *Parp1^+/A^* mice lymphocyte and myeloid development, as well as naïve B lymphocytes CSR efficiency (D). The Student’s T-test was applied to calculate the P value. ns: p>0.05.

### *Parp1*-E988A forms persistent foci at micro-radiation sites without exchange defects

To ascertain whether E988A extends the appearance of PARP1 at DNA damage sites, we measured the kinetics of foci formation by GFP-tagged PARP1 following 405nm micro-irradiation in PARP1 knockout cells (11). We chose the PARP1 knockout cells to avoid the endogenous PARP1 from interfering with the dynamics of the GFP-tagged PARP1. Both GFP-PARP1-E988A and GFP-PARP1-WT proteins rapidly formed foci within 1 minute of irradiation, suggesting the E988A mutation does not affect DNA binding by PARP1 (Fig. 4A and 4B). As expected, cells expressing GFP-PARP1-E988A failed to form bright PAR-dependent XRCC1 foci, consistent with the lack of PARylation activity (Fig. 4A, 4C and *Fig. S2B*). However, unlike GFP-PARP1-WT foci, which dissolved within ∼10 minutes, at least 35% of GFP-PARP1-E988A foci persisted for 20 minutes and 25% lasted to 30 minutes (Fig. 4A and 4D). The presence of PARP1 E988A at the DNA ends in later time points could physically block the recruitment of other DNA repair factors, including PARP2 (see below).Persistent PARP1 foci could be caused by allosteric trapping of the same PARP1 molecule at the DNA breaks or by continuous recruitment of different PARP1 molecules to the breaks due to the lack of repair (*e.g.*, reduced PAR dependent recruitment of the XRCC1-LIG3 complex) (11). To distinguish these two possibilities, we measured florescence recovery after photo-bleach (FRAP). Upon photobleaching, GFP-PARP1-E988A foci recovered as efficiently as the WT control (11) (Fig. 4E and 4F), suggesting that E988A mutation does NOT affect the exchange of PARP1 at the DNA damage foci and the persistency of PARP1-E988A foci is likely caused by continuous recruitment of different PARP1-E988A proteins to the unrepaired breaks. This is similar to what we and others have reported for clinical PARPi-treated PARP1-WT (11). Thus, this data suggests that inactive PARP1 (E988A) can occupy the DNA breaks for an extended time, where it blocks other repair proteins from accessing the ends.

**Figure 4.**
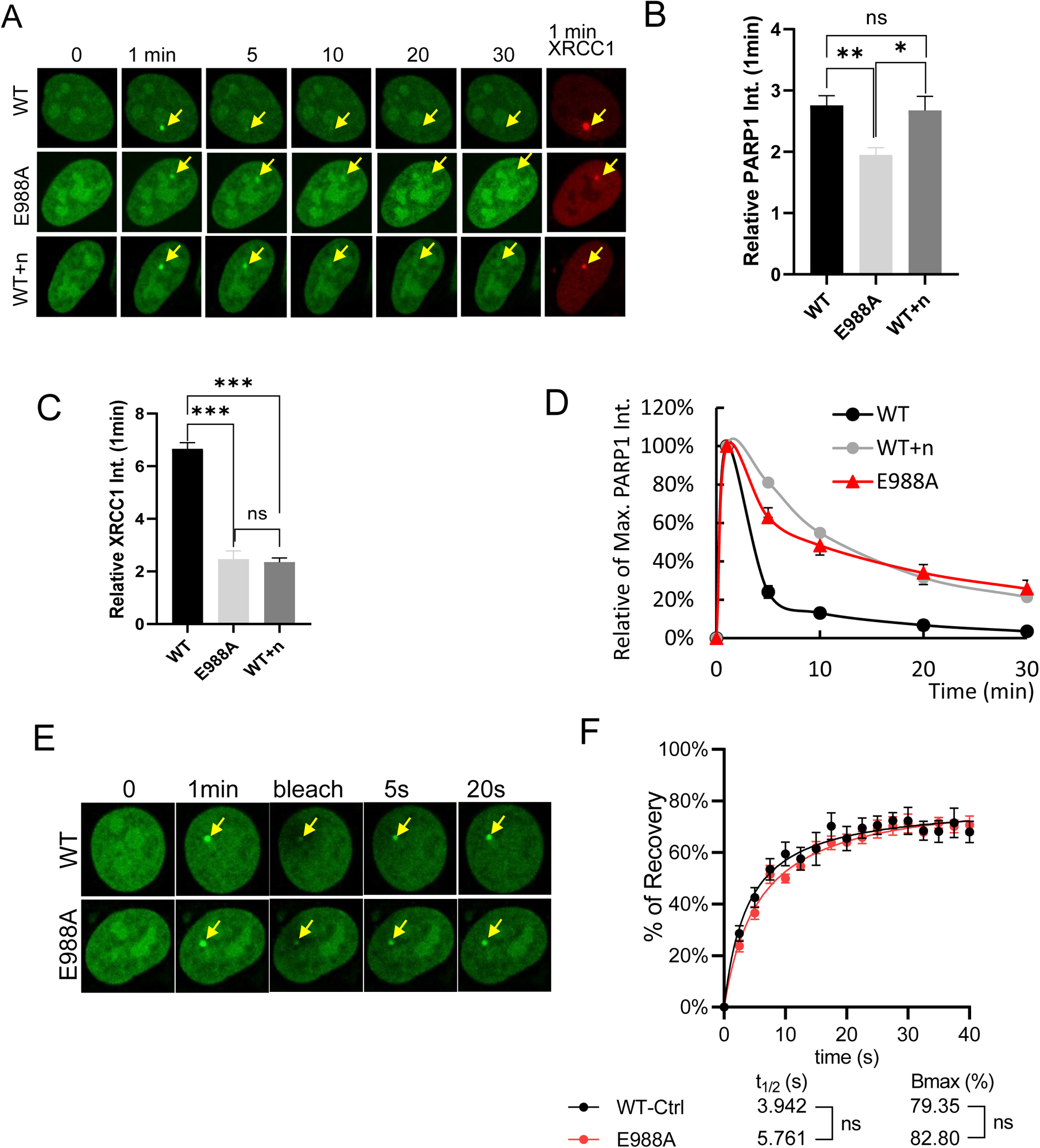
Inactive Parp1 forms persistent DNA damage-induced foci without exchange defects. (A-D) Representative images (A) (n = niraparib), 1min PARP1 foci intensity (B), 1-minute intensity of RFP tagged XRCC1 (C), and normalized kinetics (D) of GFP-tagged E988A-PARP1 or wild type PARP1 in PARP1 knock out U2OS cells with or without 1μM niraparib after laser micro-irradiation. ns: P>0.05, *: P<0.05, **: P<0.01, and ***: P<0.001. (E and F) Representative FRAP images (E) and recovery curves (F) of E988A-PARP1 and wild-type PARP1.

### Isolated *Parp1^+/A^* cells support significant DNA damage-induced PARylation

Next, we isolated *Parp1^+/A^* and *Parp1^-/-^* cells and confirmed by genotyping (*Fig. S2C-D* and Tab. S1) to understand how inactive Parp1 dominant negatively blocks embryonic development. We also validated that the A allele is expressed in the *Parp1^+/A^* cells via RT-PCR. The nucleotides encoding Glutamine 988 reside at the junction between Exon 22 and 23 (*Fig. S2E*). The targeting created a new MluI digestion site in the cDNA (but not in genomic DNA). MluI digestion of RT-PCR products derived from *Parp1^+/A^* cells detected two bands of equal intensities, indicating the A allele is expressed at levels comparable to the WT allele (*Fig. S2E*). Correspondingly, sequencing of the cDNA derived from *Parp1^+/A^* cells also revealed mixed cDNA at the A->C mutation (leading to Glu to Ala) and silent mutation T->C (Asn-> Asn) (*Fig. S2F*). Together these data confirmed that the Parp1^A^ allele is expressed at a comparable level to the WT allele in the *Parp1^+/A^* cells. In previous studies, we showed that the majority of PARP2 is recruited to DNA damage sites by the PAR chain generated by PARP1 and that the presence of PARP1 protein prevents DNA dependent recruitment and activation of PARP2 (24). Given loss of both PARP1 and PARP2 led to embryonic lethality, we hypothesized that the presence of inactive PARP1 might simply abolish the PARylation by both PARP1 and PARP2 by blocking the remaining PARP1 and PARP2 activation. To test this, we first measured PARylation dependent recruitment of XRCC1 to DNA damage foci in *Parp1^+/A^* cells. The intensity of the RFP-XRCC1 foci in *Parp1^+/A^* cells was higher than *Parp1^-/-^* cells while also lower than *Parp1^+/+^* cells (Fig. 5A). Next, we measured pan-ADP-ribosylation using Western Blot. Both baseline and damage induced auto-ADP-ribosylation of PARP1 and histones are comparable in *Parp1^+/A^* and control *Parp1^+/+^* cells and significantly higher than that of *Parp1^-/-^* cells (Fig. 5B-C). This is consistent with the notion that the E988A mutation preserves some mono-ADP-ribosylation (MARylation) activity (11). Consistent with Parp1 (regardless of activity) blocking PARP2 activation, Parp2 auto-ADP-ribosylation is highest in *Parp1^-/-^* cells, and lower in both *Parp1^+/+^* and *Parp1^+/A^* cells. Next, we used an antibody specific for PARylation (cannot detect MARylation). We found that H_2_O_2_-induced auto-PARylation in *Parp1^+/A^* cells is comparable with that of *Parp1^+/-^* cells, and much higher than that of *Parp1^-/-^* cells (Fig. 5D). This result is consistent with the intra-molecular auto-PARylation mode of PARP1. In both *Parp1^+/A^* and *Parp1^+/-^* cells only one copy of the PARP1 can be auto-PARylated. Collectively, the result suggests that the embryonic lethality of *Parp1^+/A^* mice cannot be explained by the lack of overall ADP-ribosylation or PARylation. Instead, the continuous recruitment and the presence of inactive Parp1 protein at DNA damage lesions might physically block DNA repair by preventing other proteins (e.g., PARP2) from accessing the ends.

**Figure 5.**
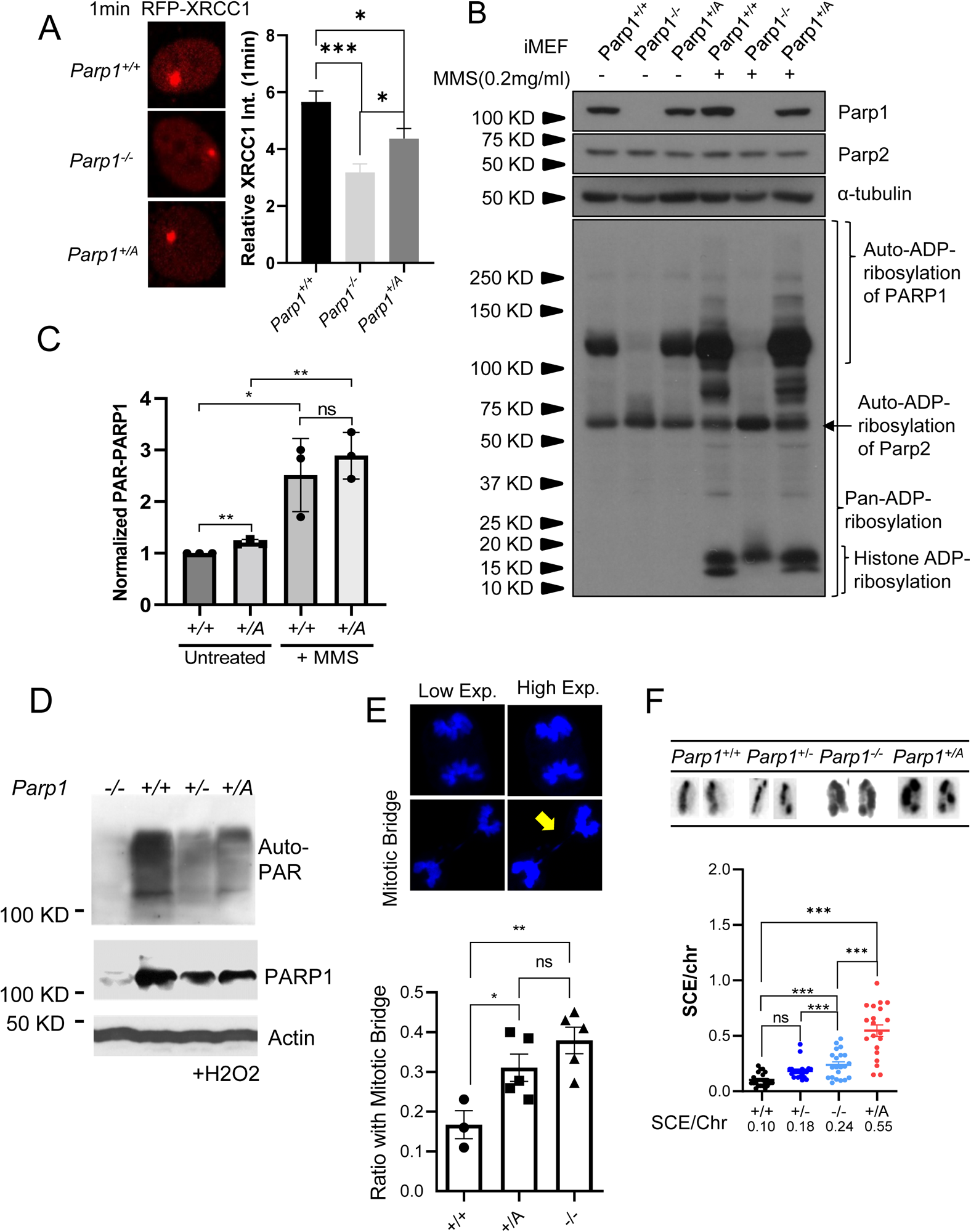
PARP1 E988A causes increased SCE in a dominant negative manner without ablating DNA damage-induced PARylation in *Parp1^+/A^* cells. (A) Representative images and 1 min intensity of RFP tagged XRCC1 foci in *Parp1^+/+^, Parp1^-/-^,* and *Parp1^+/A^* immortalized MEF after 405nm laser micro-irradiation. (B-C) Representative Western blot (B) and quantification of pan-AND-ribosylated PARP1 and PARP2 in *Parp1^+/+^, Parp1^-/-^,* and *Parp1^+/A^* immortalized MEF after 0.2mg/ml of MMS treatment (C). *: P<0.05 and **: P<0.01, P-value was calculated by Student’s T-test. (D) Western blot of PAR in H_2_O_2_ stimulated *Parp1^-/-^, Parp1^+/+^, Parp1^+/-^,* and *Parp1^+/A^* immortalized MEFs. (E) Representative images showing normal mitotic cells (top) or cells with a mitotic bridge (bottom) and frequency of mitotic bridges in *Parp1^+/+^, Parp1^-/-^,* and *Parp1^+/A^*immortalized MEF. (F) Representative images and statistics of sister chromatid exchange (SCE) rate of *Parp1^+/+^, Parp1^+/-^, Parp1^-/-^,* and *Parp1^+/A^* ES cells. The P value was calculated based on one-way ANOVA. ***: P<0.001

### Inactivated PARP1 protein selectively blocks single-strand breaks repair and Topo-II lesions

DNA strand breaks, both single and double-strand breaks, recruit and activate PARP1. To understand the types of DNA repair events blocked by *Parp1^A^*, we isolated *Parp1^+/A^* and *Parp1^-/-^* embryonic stem (ES) cells and confirmed via PCR that they do not have the *Parp1^AN^* allele (*Fig. S2C-D*). ES cells were chosen because their inherent checkpoint defects would allow us to visualize chromosomal instability in metaphase preparations easily. Mitotic bridge formation was increased in both *Parp1^+/A^* and *Parp1^-/-^* cells to similar levels (Fig.5E). Meanwhile, in comparison to the *Parp1^+/+^* cells, the frequency of sister-chromatid exchanges (SCEs) per chromosome increased ∼2.4 fold in *Parp1^-/-^* cells, and ∼ 5.5 fold in *Parp1^+/A^* cells(Fig. 5F). Despite the genomic instability, SV40-large and small T antigen immortalized *Parp1^+/A^* and *Parp1^-/-^* murine embryonic fibroblasts (iMEFs) proliferated at similar levels as the matched *Parp1^+/+^* control (5) (Fig. 6A).

**Figure 6.**
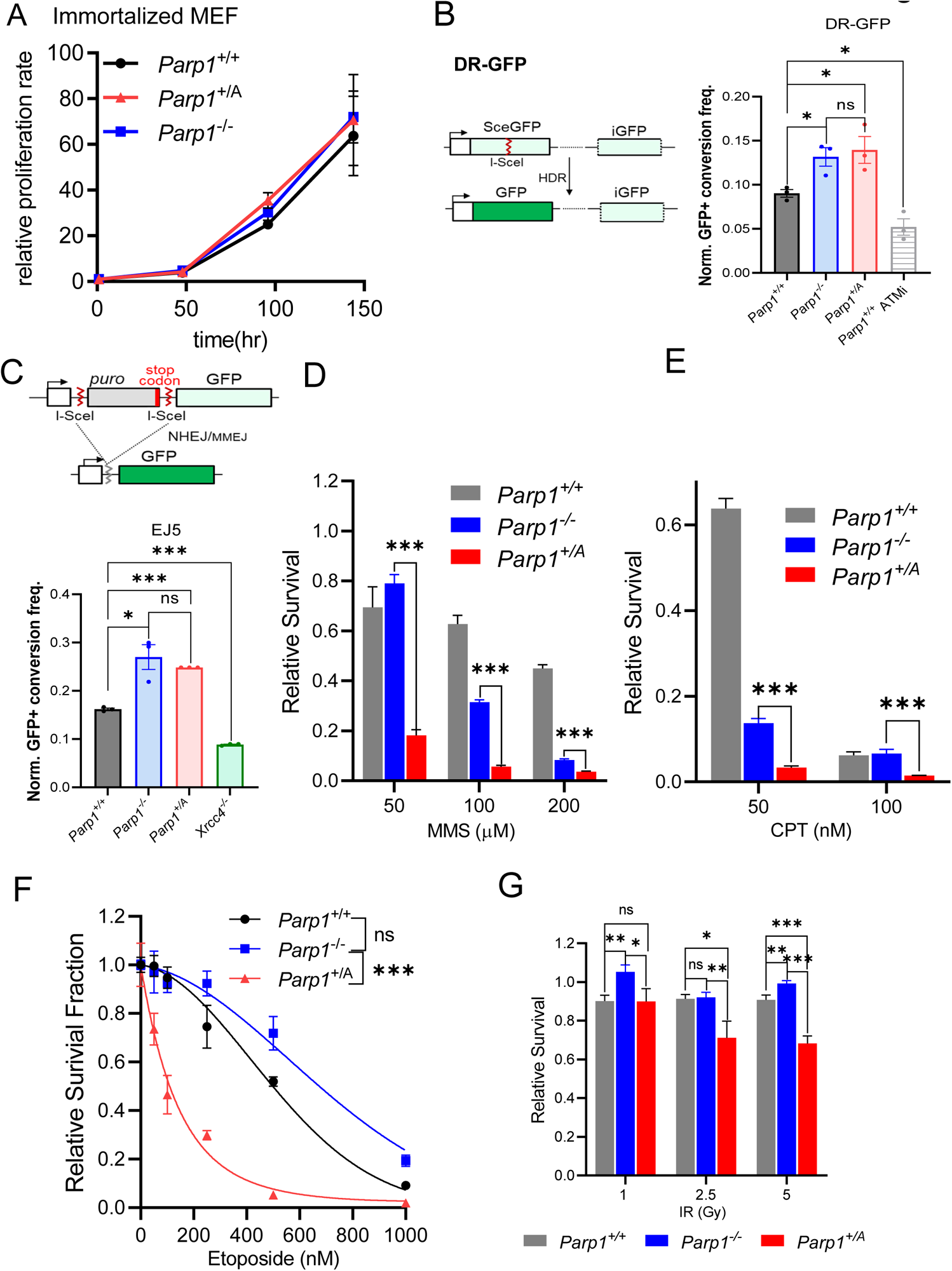
PARP1 E988A dominant negatively inhibits selective DNA repair pathways without ablating HR or NHEJ repair. (A) The proliferation rate of *Parp1^+/+^, Parp1^-/-^, and Parp1^+/A^* immortalized MEFs. (B) Sketch diagram and HR efficiency measured by the frequency of GFP+ cells in *Parp1^+/+^, Parp1^-/-^,* and *Parp1^+/A^* immortalized MEF with *Parp1^+/+^*+ ATMi (KU-55933 15uM, Selleckhem, S1092) control using DR-GFP reporter. Student’s T-test calculated P-value, ns: P>0.05 and *: P<0.05. (C) Sketch diagram and NHEJ efficiency measured by the frequency of GFP+ cells in *Parp1^+/+^, Parp1^-/-^,* and *Parp1^+/A^* immortalized MEF with *Xrcc4^-/-^*MEF control using EJ5 reporter. P-value was calculated by Student’s T-test, ns: P>0.05, *: P<0.05, and ***: P<0.001. (D-G) Sensitivity of *Parp1^+/+^, Parp1^-/-^,* and *Parp1^+/A^* immortalized MEFs to MMS (D), CPT (E), etoposide (F), and IR(G). P-value was calculated by Student’s T-test and extra sum-of-square F test of the dose-response model. All dots and error bars represent means and standard errors. ns: P>0.05, *: P<0.05, **: P<0.01, and ***: P<0.001.

Using this genotype and expression validated *Parp1^+/A^* iMEF, we measured HR and NHEJ (and microhomology mediated end-joining -MMEJ) using the DR-GFP and EJ5 reporters, respectively. After the reporter has been integrated into the *Parp1^+/A^* and control iMEF, transient expression of I-SceI induces GFP conversion via HR in DR-GFP reporter containing cells (25) and via NHEJ, and to a lesser extent MMEJ, in EJ5 containing cells (26). Consistent with normal meiosis in the young *Parp1^+/A^* F1 chimeras, *Parp1^+/A^* and *Parp1^-/-^* iMEFs support DR-GFP conversion at levels comparable to or even higher than the *Parp1^+/+^* iMEFs (Fig. 6B). ATM inhibitor treatment significantly reduced DR-GFP conversion in *Parp1^+/+^* cells, serving as a control (Fig. 6B)(27–29). In parallel, EJ5 conversion is also very efficient in *Parp1^+/A^* and *Parp1^-/-^* iMEFs (Fig. 6C). The elevated levels of DR-GFP in *Parp1^+/A^* and *Parp1^-/-^* iMEFs are not due to high copies of the substrates in *Parp1^+/A^* and *Parp1^-/-^* iMEFs (Fig. S3A-B) but might reflect the increased SCE in *Parp1^+/A^* and *Parp1^-/-^* cells (Fig. 5F). Together with the normal lymphocyte development and meiosis in the young F1 *Parp1^+/A^* mice, the results suggest that inactive PARP1 did not abrogate HR or NHEJ.

To understand what other type of DNA repair was blocked by inactive PARP1, we measured DNA damage sensitivity using *Parp1^+/A^* and control iMEF. Even in comparison to *Parp1^-/-^* iMEFs, *Parp1^+/A^* iMEFs were much more sensitive to the alkylating agent methyl-methanesulfonate (MMS) (Fig. 6D and S3C), the Topoisomerase I inhibitor camptothecin (CPT) (Fig. 6E and S3D) and, perhaps unexpectedly, the Topoisomerase II inhibitor etoposide (Fig. 6F). *Parp1^-/-^* cells are also sensitive to MMS and CPT, albeit less than their *Parp1^+/A^* counterpart, consistent with a role of PAR and PAR-mediated recruitment of XRCC1 in base excision repair and Top1-poison removal (Fig. 6D and 6E). In this context, XRCC1 deficiency has also been linked with MMS and CPT hypersensitivity (2). But etoposide sensitivity has not been reported in XRCC1-deficient cells. The hypersensitivity of *Parp1^+/A^* cells to Topo-II block and the normal lymphocyte development and spermatogenesis in young F1 *Parp1^+/A^* chimera mice, together with their HR and NHEJ/MMEJ proficiency measured by the DR-GFP and EJ5 reporters (Fig. 6B-C), suggest inactive PARP1, but not lack of PARP1, might block the processing of end-blocked lesions, especially during replication, likely independent of NHEJ or HR. Consistent with this notion, *Parp1^+/A^* and *Parp1^-/-^* iMEF displayed consistent and comparable levels of hypersensitivity to radiation (Fig.6G and S3E).

## Discussion

Taken together, the early embryonic lethality of *Parp1^+/A^* mice provides the first evidence for substantial toxicity of catalytically-inactive PARP1 protein in otherwise normal tissues. This is in sharp contrast to the normal development of *Parp1^-/-^* mice. In this context, *Parp1^+/A^* cells share many features with PARPi-treated normal cells (18) and XRCC1-deficient cells(2, 3), including BER deficiency, increased SCE, and persistent PARP1 foci without allosteric locking(11, 14). Conversely, unlike the PARPi-treated cells, the overall ADP-ribosylation activity in the *Parp1^+/A^* cells closely resembles the *Parp1^+/+^* cells, and is much higher than in *Parp1^-/-^* cells (Fig.5B-C). The damage-induced auto-PARylation of PARP1 in *Parp1^+/A^* cells is similar to *Parp1^+/-^* cells (Fig.5D), consistent with both lines having only one copy of active PARP1 that can auto-PARylate itself. Loss of PARP1 leads to resistance to PARP inhibitors (11, 30). Together these data support a model in which the presence of inactive PARP1, but not the lack of PARP1-mediated PARylation in general, underlies the normal tissue toxicity of PARP inhibition. Moreover, it suggests that if half of the PARP1 is inactive, it is sufficient to cause major normal tissue toxicity. Given no small molecule inhibitors are present in the *Parp1^+/A^* cells and the efficient exchange of PARP1-E988A at the micro-irradiation sites, our results also suggest that the small molecule induced reverse allosteric locking of PARP1, although may be important for inhibitor function, are not essential for normal tissue toxicity. It is the presence of inactive PARP1 with high affinity DNA binding that is toxic. In this context, the kinetics of PARylation, thus enzymatic activity might important for preventing persistent PARP1 foci (31). HPF1 was recently identified as an essential modulator for PARP1 activity on Serine (32). Lack of HPF1 and similar modulators also cause persistent PARP1 foci and toxicity, offering new strategies for targeting PARP1.

How does the inactive PARP1 differ from PARP1 deletion? MMS induced PARP2 auto-ADP-ribosylation increased in *Parp1^-/-^* cells, but not in *Parp1^+/A^* cells (Fig. 5B-C), suggesting the presence of inactive Parp1 blocks PARP2 activation and auto-modification. It is possible PARP2 is only one of the many repair proteins that is blocked by inactive PARP1. As a result, while both *Parp1^+/A^* cells and *Parp1^-/-^* cells are hypersensitive to MMS and Topo I inhibitors, the hypersensitivity to Topo II inhibitors seems unique to the presence of inactive PARP1 and has not been found with PARP1 or XRCC1 deletion (2, 3). Among the genotoxic anti-cancer therapies, alkylating agents and Topo II inhibitors carry a particularly high risk for therapy-induced MDS/AML (t-MDS/AML) and clonal hematopoiesis. Thus, the hypersensitivity of *Parp1^+/A^* cells to these agents might explain the severe hematological toxicity and t-MDS/AML associated with PARP inhibitors(17, 19), a hypothesis that can be tested by somatic inactivation models.

The *Parp1^+/A^* mouse model described here provides the first evidence for severe toxicity of the inactive PARP1 protein during embryonic development *in vivo*. PARP1 resembles the ATM, ATR, and DNA-PK kinases, which also exhibit more severe phenotypes when enzymatically impaired than when deleted entirely (33). They all share a common feature. All are recruited to and activated at the site of DNA damage. The activation is coupled with a rapid exchange of the proteins at the DNA damage site. The presence of the inactive protein not only blocks the further activation of the signaling cascade, but also occupies the DNA lesion, where it blocks the activation of other sensors or DNA repair itself. This model is consistent with the fact that no “inactivation” mechanism for these DNA damage response factors exists. These insights should guide the development of new therapeutic agents that circumvent the normal tissue toxicities associated with current PARP1, ATM, and ATR inhibitors.

## Materials and Methods Mice

The 5’ and 3’ arms of the Parp1^E988A^ targeting construct were amplified from 129/sv mouse embryonic stem (ES) cells’ genomic DNA, mutated in exon22 encoding E988A, and cloned into the pEMC-neo targeting vector with a PGK-neo-resistance (neo-R) cassette flanked by FRTs. Targeted ES clones were screened by PCR first and confirmed via Southern blotting (HindIII and KpnI digestion with both 5’ and 3’ probes) (*Fig. 1B* and Tab. 1). The successfully targeted ES clones were injected for germline transmission. The *Parp1*^+/AN^ chimeras with the neo-R cassette were bred with Rosa26a^FLIP/FLIP^ (Jax Strain No. 003946 in 129Sv) mice to generate the F1 *Parp1*^+/A^ mice. The presence of the NeoR in Parp1^AN^ allele interferes with the expression of Parp1, resulting in no or little Parp1 expression. The p-value of birth rates was calculated by χ^2^ test. Genotypings were carried out by PCR with 230bp product for *Parp1^+^*, 315bp for *Parp1^A^*, and 440bp for *Parp1*^AN^ in the triplex PCR, and 340bp for only AN (Fig. 2F, 2G and *Fig. S1C, S1D, S2C, S2D, and Tab. 1*). All animal work was carried out with the guidelines established by the Institutional Animal Care and Use Committee (IACUC) at Columbia University Medical Center.

### Western blot antibodies

Primary antibodies used in this study for Western blot include: anti-PARP1 antibody (Cell Signaling Technology, 9542, 1:2000), anti-PARP2 antibody (Active Motif, 39044, 1:2000), anti-α-Tubulin antibody (Sigma, CP06, 1:1000), anti-pan-ADP-ribose binding reagent (Sigma, MABE1016, 1:1000), anti-β-actin (Sigma, A5441, 1:10000), and anti-PAR antibody (R&D, 4335-MC-100, 1:1000).

### Mouse hematopoietic cells and lymphocytes analysis

Mouse blood cell and lymphocyte development were analyzed by flow cytometry(34, 35). B lymphocyte development was conducted by staining with cocktails including FITC anti-mouse CD43 (Biolegend, 553270), PE goat anti-mouse IgM (Southern Biotech, 1020-09), PE-cyanine5 anti-Hu/Mo CD45R (B220) (eBioScience, 15-0452-83), and APC anti-mouse TER119 (Biolegend, 116212). T lymphocyte development was measured on thymocytes using cocktail including PE rat anti-mouse CD4 (Biolegend, 557308), FITC anti-mouse CD8a (Biolegend, 100706), PE/Cy5 anti-mouse CD3e (eBioscience, 15-0031-83), and APC anti-mouse TCRβ (BD Pharmingen, 553174). Myeloid cells were measured in bone marrow cells and splenocytes using cocktail including FITC anti-mouse CD11b (BD Pharmingen, 553310), PE rat anti-mouse CD19 (BD Pharmingen, 557399), PE/Cy5 anti-mouse CD3e (eBioscience, 15-0031-83), and APC anti-mouse Ly6G/Ly6C(Gr-1) (Biolegend, 108412). Purified splenocytes CSR was stained with PE-cyanine5 anti-Hu/Mo CD45R (B220) and FITC rat anti-mouse IgG1(BD Pharmingen, 553443). All antibodies were diluted according to manufacturer’s protocol. The flow cytometry data were collected on either an LSR II (BD), or on an Attune NxT (Invitrogen) flow cytometer. All flow cytometry data were analyzed using FlowJo V10.

### Cell line derivation and sensitivity assay

The *Parp1*^+/+^, *Parp1*^-/-^, *Parp1^+/-^*, and F1 *Parp1^+/A^* MEFs were derived from E13.5–E14.5 embryos and functionally and genotyping verified (Fig. S2C-D). The *Parp1*^+/+^, *Parp1*^-/-^, *Parp1*^+/-^ and *Parp1*^+/A^ ES cells were derived at the HICCC transgenic mouse core facility and cultured on irradiated (30Gy) fibroblast feeders as detailed before(36, 37). For the survival assay, ∼1000 cells/well were plated into a 96-well plate. Cells were irradiated by ionizing radiation (IR, 1, 2.5, or 5Gy) 24hr after plating. Genotoxins were added 24hrs after initial plating at the concentration indicated in the figures - camptothecin (CPT, Sigma, 208925) 50, and 100 nM; etoposide (Sigma, E1383) 50, 100, 250, 500, and 1000 nM; and methyl-methanesulfonate (MMS, Sigma, 129925) 50, 100, and 200 μM. The CyQUANT™ Cell Proliferation Assay kit (Invitrogen, C7026) was used to measure cell numbers after 7 days. The survival curves were generated with GraphPad Prism V8.0 using the nonlinear regression linear quadratic cell death model and P-value was calculated based on coefficient of killing.

### Adult male mouse epididymal sperm count and sperm DNA preparation

Mice sperm counts and histology analyses of the adult testis (<8 weeks) were carried out as detailed before(38). Testes were fixed in Bouin’s fixative (ICCA, 1120-16) for Periodic Acid Schiff (PAS) staining. Cauda epididymis was minced in PBS and incubated at 32°C for 20 min to let all sperms swim out. Sperm suspension was filtered through a 70-μm nylon cell strainer and counted under a microscope using a hematocytometer. Sperm mobility was determined as: Grade a: immotile, b: non-progressive, c: slow progressive, or d: rapid progressive.

### Live cell imaging

Live cell imaging and FRAP were conducted as previously described with minor revisions (10, 11). Plasmids encoding GFP-tagged WT or E988A PARP1, together with RFP-tagged XRCC1 (a generous gift from Dr. Li Lan at MGH), were transfected into PARP1 knockout U2OS cells (11) using Lipofectamine 2000 (Invitrogen, 11668019). Live-cell images were acquired via the NIS Element High Content Analysis software (Nikon Inc.) at different times after damage (inflicted by 405 nm laser without priming) and every 2.5s after photobleaching (via a 488 nm laser) at 1 min after initial damage for FRAP. P-values of exchange curves were calculated based on extra sum-of-square F test of the dose-response model. All Dots and error bars represent means and standard errors. All images’ analyses were carried out with Fiji ImageJ software.

### HR and NHEJ reporter assay

The DR-GFP, EJ5 and I-Scel plasmids were generous gifts from Dr. Jeremy Stark at the City of Hope and the assays were carried out as previously detailed (26) with minor modifications. Briefly, the DR-GFP plasmid contains an upstream SceGFP cassette in which the GFP expression was interrupted by an I-Scel recognition site, and a downstream GFP template truncated at both termini. After I-Scel induced DSB, a successful HR repair using the downstream iGFP as template would convert the SceGFP cassette into an expressing GFP cassette. The EJ5 plasmid consists of a GFP cassette downstream of a puromycin ORF flanked by two I-Scel sites following a single promoter, in which the stop codon in the puromycin ORF prevents GFP expression. Successful NHEJ repair following I-Scel digestion restores GFP expression. Both the DR-GFP and EJ5 plasmids were linearized and electroporated (BioRad Gene Pulser Xcell™ system, 800V, 10uF, 300ohm, 0.4cm cuvettes) into immortalized *Parp1*^+/+^, *Parp1*^-/-^, F1 *Parp1^+/A^* and *Xrcc4^-/-^* MEFs separately. After 6 days of puromycin selection and expansion, I-Scel plasmid was transiently transfected into the stable cell lines using lipofectamine 2000 following manufacturer’s protocol. Frequency of GFP expressing cells was measured by flow cytometry at D3 after I-Scel transfection, and HR and NHEJ efficiency was calculated by normalizing GFP+ population to the efficiency of GFP expressing vector transfection control to elucidate the effect of transfection efficiency. All reporter assays were repeated for additional 2 times and all flow cytometry data were analyzed using FlowJo V10.

## Acknowledgments

We thank Dr. Chyuan-Sheng (Victor) Lin for helping with germline injection, Dr. Theresa Swayne for helping with confocal analyses, and Dr. Jeremy Stark for his great help and discussion about the DR-GFP and EJ5 reporter assays. This work was partly supported by NIH R01 CA226852, CA271595, CA275184 to SZ, HD085904 to MMS, and 1P01CA174653 to SZ and RB. SZ was a scholar of the Leukemia & Lymphoma Society (LLS). This research used the flow cytometry, molecular cytogenetic, transgenic, and confocal and specialized microscopy shared resources funded through the NIH/NCI Cancer Center Support Grant P30CA013696 to the HICCC of Columbia University.

---

**SI Appendix Figure 1.**
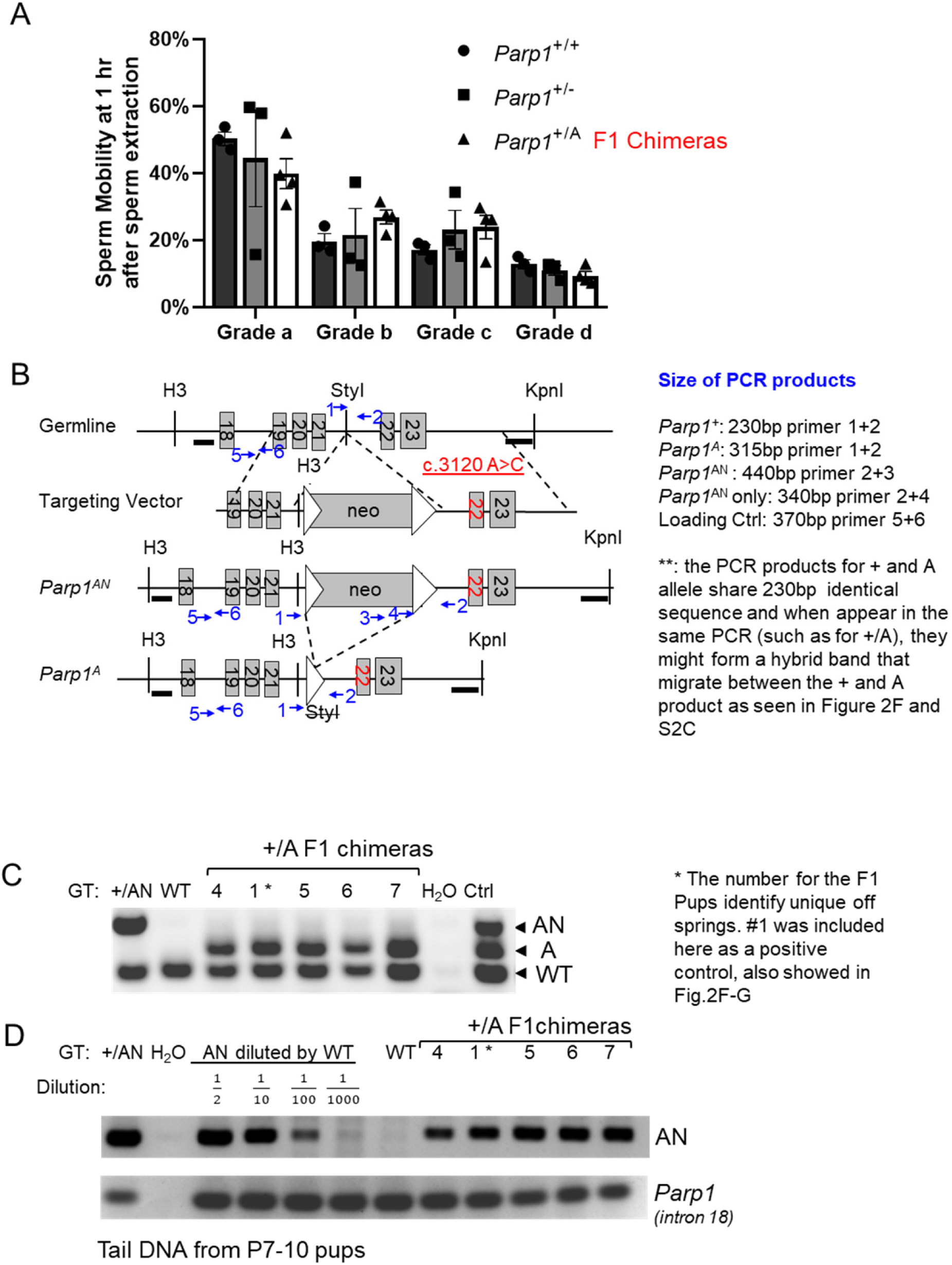
Spermatogenesis, genotype primers, and the genotyping result of additional *F1 Parp1^+/A^* mice. (A) Mobility of epididymis sperm of *Parp1^+/+^*, *Parp1^+/-^*, and *Parp1^+/A^* 8–24-week F1 chimeric adult males. One-way ANOVA calculated the P-value. No significant difference was identified in each group. (B) Position and product size of PCR primers for triplex PCR and AN-specific PCR marked on the targeting diagram. Blue arrows indicate primer positions. (C-D) Standard triplex tail PCR (C) and targeted AN PCR (D) of other F1 *Parp1^+/A^* mice and controls. The numbers denoted the ID of individual pups. Pup 1 was used in Fig. 2 and also here as the control.

**SI Appendix Figure 2.**
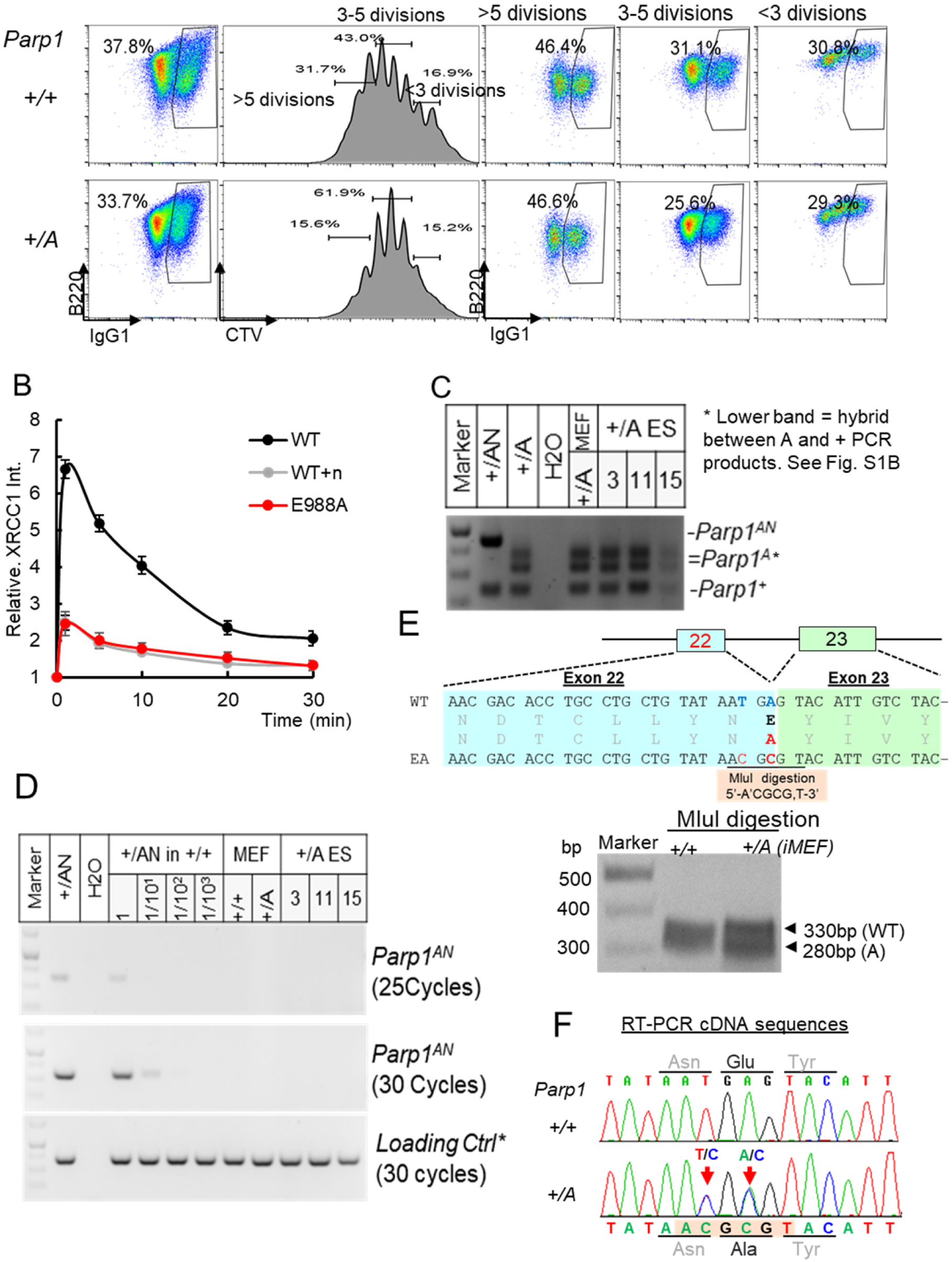
Lymphocyte development, additional cell biology analyses, and PCR verification for the genotypes. (A) Representative flow cytometry analyses of *Parp1^+/A^* mice CD43 negative splenocytes CSR with CTV staining showing passage progression. (B) Normalized kinetics curve of RFP tagged XRCC1 in GFP tagged E988A-PARP1 or wild type PARP1 expressing U2OS cells with or without 1μM niraparib. (C-D) Triplex PCR (C) and Targeted AN-PCR (D) identified genotype purity of *Parp1^+/A^* MEF, ES cell, and controls used in sensitivity assay, mitotic bridge observation, HR and NHEJ reporter assay. (E-F) Diagram of *Parp1^A^* cDNA identification and verification of *Parp1^A^* allele expression by RT-PCR with MluI digestion(E) and Sanger sequencing (F).

**SI Appendix Figure 3.**
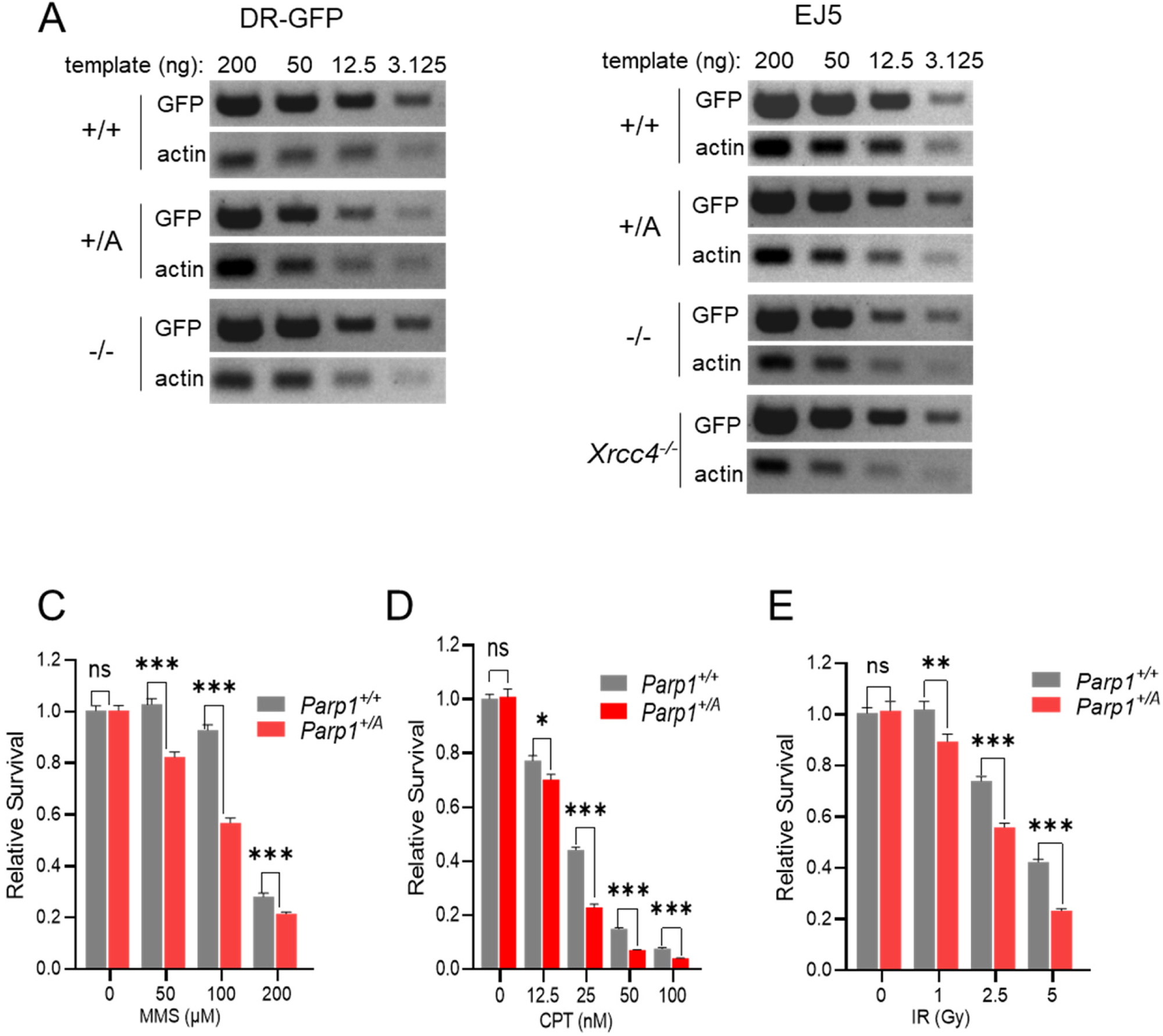
PARP1 E988A expression causes additional genomic instability. (A-B) Dilution PCR with GFP in DR-GFP (A) and EJ5 reporter (B) and controlled actin primers showed similar levels of substrates (GFP): actin copy ratio in *Parp1^+/+^, Parp1^-/-^, Parp1^+/A^,* and *Xrcc4^-/-^* immortalized MEFs. (C-E) Repeats of sensitivity assay of *Parp1^+/+^* and *Parp1^+/A^* to MMS (C), CPT (D), and ionizing radiation(E). All dots and error bars represent means and standard errors. ns: P>0.05, *: P<0.05, **: P<0.01, ***: P<0.001. P values were calculated by multiple student’s t-tests.

**SI Appendix Table 1.**
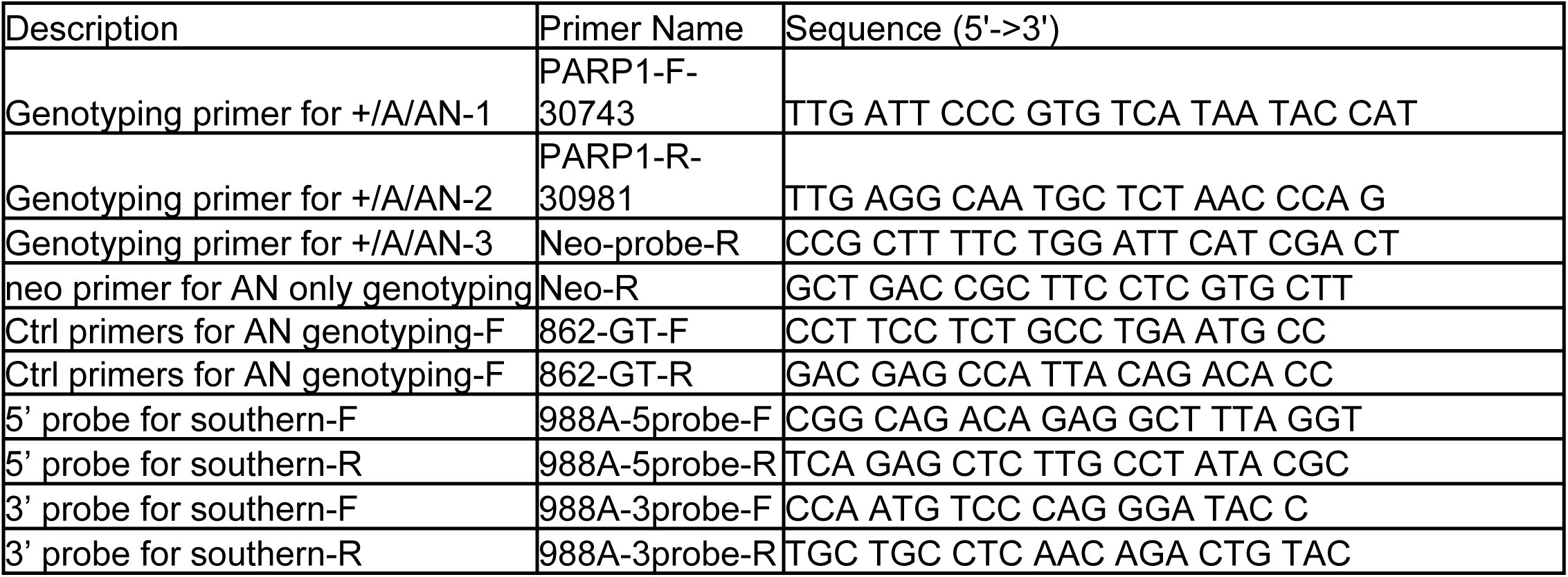
Primers used in this study

